# Hidden biodiversity in Neotropical streams: DNA barcoding uncovers high endemicity of freshwater macroinvertebrates at small spatial scales

**DOI:** 10.1101/2020.03.31.018457

**Authors:** Luis F. De León, Aydeé Cornejo, Ronnie G. Gavilán, Celestino Aguilar

## Abstract

Aquatic macroinvertebrates play a crucial role in freshwater ecosystems, but their diversity remains poorly known, particularly in the tropics. This “taxonomic void” represents limits our understanding of biodiversity patterns and processes in freshwater ecosystems, and the scale at which they operate. We used DNA barcoding to estimate lineage diversity (and the diversity of unique haplotypes) in 224 specimens of freshwater macroinvertebrates at a small spatial scale within the Panama Canal Watershed (PCW). In addition, we compiled available barcoding data to assess macroinvertebrate diversity at a broader spatial scale spanning the Isthmus of Panama. Consistently across two species delimitation algorithms (i.e., ABGD and GMYC), we found high lineage diversity within the PCW, with ~ 100-106 molecular operational taxonomic units (MOTUs at 2% sequence divergence) across 168 unique haplotypes. We also found a high lineage diversity along the Isthmus of Panama, but this diversity peaked within the PCW. However, our rarefaction/extrapolation approach showed that this diversity remains under sampled. As expected, these results indicate that the diversity of Neotropical freshwater macroinvertebrates is higher than previously thought, with the possibility of high endemicity even at narrow spatial scales. Geographic isolation is likely a main factor shaping these patterns of diversity. However, local disturbances such as the rupture of the continental divide due to the construction of the Panama Canal might be reshaping these patterns of diversity at a local scale. Although further work is needed to better understand the processes driving diversification in freshwater macroinvertebrates, we suggest that Neotropical streams represent continental islands of diversity. Understanding these islands of diversity is crucial in the face of increasing human disturbance.

## Introduction

Aquatic macroinvertebrates are a fundamental component of Neotropical freshwater environments. They mediate important processes such as food web dynamics, energy flow and nutrient cycling, and therefore play a central role in sustaining the biodiversity and functioning of freshwater ecosystems [1–3]. However, the diversity of Neotropical freshwater macroinvertebrates remains poorly described, and even less is known about the processes that drive their diversity, and the scale at which they operate [4]. For instance, despite considerable efforts by local taxonomists [e.g., 5–8], only a limited number of species are recognized (e.g., [9,10], and most of the published literature use genus and family as a standard taxonomic unit for Neotropical macroinvertebrates [9,11–15]. This is partially due to the complexity of these communities, which are often composed of multiple life-stages existing at the interface between the terrestrial and aquatic environment [16,17]. Another limitation is the low accuracy and precision of traditional morphological methods, which are generally time-consuming and not necessarily applicable across taxa and experts.

This “taxonomic void” has important consequences for our general understanding of biodiversity patterns and processes, both in Neotropical environments and globally. For example, species diversity is generally expected to increase at lower latitudes [18,19]□, but no consensus has been reached for macroinvertebrates, given the current lack of taxonomic knowledge [20–23]. Within the Neotropics, our current understanding of the drivers of species diversity in benthic macroinvertebrates is also limited [24–26].

Similar to other freshwater taxa [27,28], spatial isolation is likely a major factor driving diversification in macroinvertebrates, but few studies have tested this expectation [26,29,30]. In particular, Murria et al. [26] found high frequency of unique haplotypes associated with geographical distance across watersheds in Panama. While confirmatory, these findings are not surprising, given the large geographic distance among the watersheds included in Múrria et al. [26]. However, patterns of haplotype (or lineage) diversity at smaller scales (e.g., among streams within watersheds), where dispersal and gene flow might be less restricted, have received less attention. Accordingly, at small spatial scales we might expect a reduction in molecular diversity, in contrast to larger geographic scales. To explore this issue, we use DNA-barcoding to assess patterns of lineage diversity (and the diversity of unique haplotypes) in freshwater macroinvertebrates across four streams within the Panama Canal Watershed (PCW). In addition, we compiled available barcoding data [26] to contrast macroinvertebrate diversity at a broader spatial scale, among streams along the Isthmus of Panama.

Assessing the patterns and drivers of macroinvertebrate diversity at different scales is particularly relevant, given the increasing rate of environmental degradation in Neotropical regions [31–34]. This includes alterations such as introduction of alien species [35,36], habitat degradation, water pollution, and climate change [31,34,37,38]□. In consequence, a large portion of this biodiversity risks being lost before discovery.

## Material and methods

### Study sites and sample processing

Samples were collected from four streams within the PCW (Frijolito, Frijoles, Trinidad, Indio) between April and May of 2013 (Fig 1). Frijolito (09°08’57.9” N, 79°43’53.2” W) and Frijoles (09°09O8.2” N, 79°44’05.3” W) are typical Neotropical streams separated by approximately 300 m and located inside Soberanía National Park. These streams are surrounded by dense secondary forest and present low levels of disturbance. Río Trinidad (8°58’28.50” N, 79°57’23.9” W) is located approximately 30 km West Río Frijoles in an agricultural landscape dominated by pasture, but it has abundant riparian vegetation. Rio Indio (09°12’04.1” N, 079°24’20.4” W), located 35 Km east of Frijoles, is intermediately disturbed and, is surrounded by secondary forest with dense riparian vegetation. At each site, we haphazardly collected aquatic macroinvertebrates using standard kick-netting from the two dominant habitats types (riffles and pools). Sampling effort was approximately two hours at each site. All samples were sorted in the field and immediately preserved in 95% ethanol. Sampling permit was obtained from the Autoridad Nacional del Ambiente de Panama (Permit No. SC/A-44-12).

**Fig 1.**
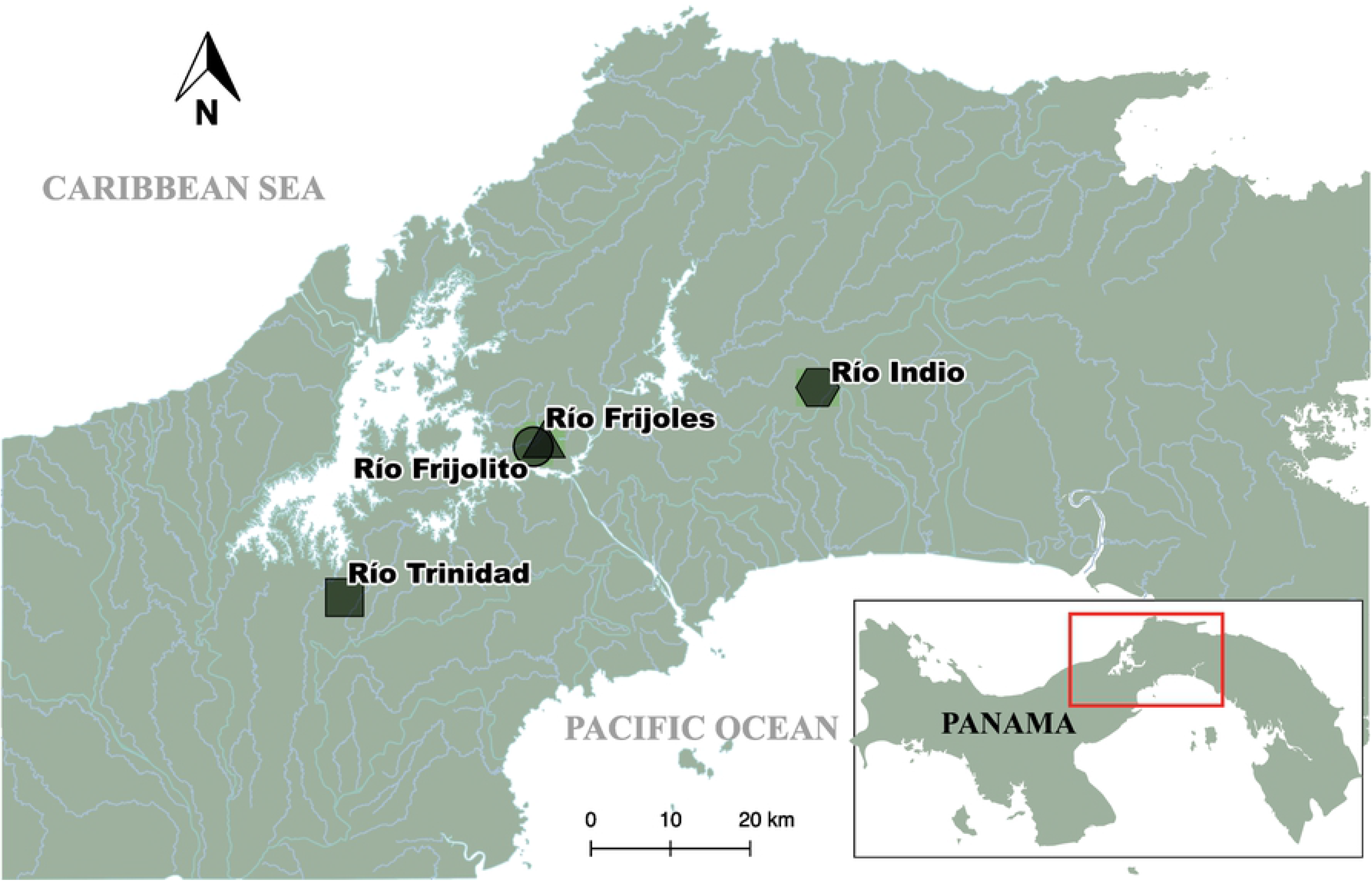
Sampling sites of macroinvertebrates in the Panama Canal Watershed.

In the laboratory, specimens were morphologically identified to the lowest possible taxonomic level (i.e., family or genus) using taxonomic keys for Neotropical macroinvertebrates [6,8,11,39]. However, given the low accuracy of morphological identification, and the fact that less than 50% of the individuals were successfully identified to species level using our barcoding data (see results), we focused our analyses and discussion on lineage rather than morphological diversity. Representative specimens have been deposited in the invertebrate collection at Colección Zoológica Dr. Eustorgio Méndez (CoZEM) at Instituto Conmemorativo Gorgas de Estudio de la Salud in Panama City (Voucher numbers: B001 — TR020).

### DNA sequencing

Tissue samples were obtained from the hind leg or part of the body of each specimen, and total DNA was extracted by using the DNeasy Blood & Tissue kit (Qiagen, CA, USA), according to the manufacturer’s instructions. A standard sequencing protocol [40] was used to amplify the full-length 658 base pair (bp) of the COI barcode region using the following primers sets: LCO1490/HCO2198 [41] and LepF1/LepR1 [42]. All PCR products were verified on a 1% agarose gel and purified with the enzymes Exonuclease I (EXO) and Shrimp Alkaline Phosphatase (SAP) [43], before sequencing using an Applied Biosystems Genetic Analyzer (ABI 3130xl, Applied Biosystems, Carlsbad, California). Sequences were aligned using Geneious V7.03 [44]. Sequence alignments were also inspected by eye to confirm overall sequence quality. Project sequences are available in GenBank (accession numbers: KX039451-KX039650, KU980966-KU981004).

### Data analysis

To confirm morphological identification for our sequenced specimens, we performed BLAST searches for publicly available sequences in GenBank. We then collected haplotype information (426 haplotypes: GenBank accession numbers KR134410-KR134835) from a previous study spanning the Isthmus of Panama [26]. After adding these sequences to our data set, we generated multiple sequence alignments with MAFFT 7.313 [45] using the L-INS-i algorithm. Then, we trimmed the sequences to the same fragment size, and compared the previously reported haplotypes with the ones encountered in our dataset. To exclude redundancies prior to phylogenetic analyses, we applied DAMBE v. 6.4.11 [46] to identify and remove duplicate haplotypes from our dataset.

We created a final dataset comprising only unique haplotypes from our study, and including three COI sequences retrieved from Genbank that were used as outgroup. The retrieved sequences were *Thermobia domestica* (GenBank NC006080), *Atelura formicaria* (GenBank NC01119) and *Tricholepidion gertschi* (GenBank NC005437).

We estimated phylogenetic relationships among taxa using maximum likelihood (ML) searches in IQ-TREE v 1.6 [47] and Bayesian inference (BI) in BEAST v 2.4.6 [48] as implemented on the CIPRES Science Gateway [49]. The best-fit model of nucleotide substitution for the dataset was selected using jModelTest 2.0 [50] based on the Bayesian Information Criterion. GTR+I+G was selected as the best-fit model for further analyses. To determine node support for the IQ-TREE we used 10000 ultrafast bootstrap [51] and 1000 Shimodaira-Hasegawa-like approximate likelihood ratio test [52] replicates. BI analysis was executed with an uncorrelated lognormal relaxed clock and coalescent prior, with the default settings of BEAUti for the remaining parameters with two runs of 2.0×10^7^ generations sampling trees every 5000 generations. Trace logs and species trees for the two runs were combined using LogCombiner v 2.4.8 [53]. We used Tracer v. 1.6 [54] to ensure that effective sample size (ESS) values for all parameters were above 200 and to determine the burn-in. Finally, output trees were summarized as maximum clade credibility (MCC) trees using mean node heights after discarding 25% of generations as burn-in using TreeAnnotator v1.8.4 [55].

We then assessed species diversity by estimating molecular operational units (MOTUs; [56]. Sequence divergence was estimated using the K2P model with 1000 bootstrap estimates in MEGA7 [57]. This a standard model that has been extensively used in barcoding studies [58,59]. MOTUs or barcode species were delimited using 2% in divergence cut-off, which has proved to be consistent for most invertebrate groups [60–62].

To confirm lineage diversity, we applied two single-locus analyses, the Bayesian General Mixed Yule Coalescent model (GMYC; [63] and the Automatic Barcode Gap Discovery (ABGD; [64] on the mitochondrial dataset. The GMYC approach uses branch lengths to determine the transition from intraspecific to interspecific relationships [63]. The ABGD algorithm allows to sort DNA sequences into “hypothetical species” based on the gaps in the distribution of intra- and inter-specific genetic divergence in a given sample [64]. Although the two approaches differ in their properties (i.e., tree branch length vs. distribution gaps), we used them to confirm the patterns of species delimitation.

To perform GMYC tree-based analyses, we used the ultrametric trees previously generated with BEAST. GMYC was performed using the single threshold parameter at the GMYC webserver (https://species.h-its.org/gmyc/). ABGD was carried out using the online version of ABGD software [64]; https://bioinfo.mnhn.fr/abi/public/abgd/abgdweb.html). Default settings were used, however, distance matrices based on K2P distance calculated in MEGA7 were used as input. All analyses were run using a relative barcoding gap width (X value) set to 1.0. Only the recursive results were used because they allowed for different gap thresholds among taxa.

To compare patterns of spatial variation in genetic diversity (MOTUs), we quantified the number of shared species among sampling sites. We also estimated Fisher’s alpha index of diversity and Whittaker’s measure of β-beta diversity. Given that these analyses might be affected by variation in sampling size, we also used rarefaction and extrapolation method [65,66] as implemented in the R package iNEXT [67]. This method allows for comparisons between sites while controlling for differences in abundance and sampling effort. For these analyses, we fit curves for the first three Hill numbers: species richness (q = 0), the exponential of Shannon entropy (“Shannon diversity”, q = 1), and the inverse Simpson concentration (“Simpson diversity”, q = 2), using individual-based abundance data. Finally, to assess lineage diversity and the diversity of unique haplotypes across watersheds spanning the Isthmus of Panama, we compiled data from Murria et al. [26]. The combined dataset contained 12 sites (8 from Múrria et al. [26], and 4 from the present study). One site (Frijolitos) was sampled during both studies, but we analyzed them separately to preserve independence between the two studies. We then applied the same rarefaction/extrapolation approach described above to generate rarefaction curves as function of the number of individuals sampled. Although there might be important methodological differences between the two studies, our objective here is to provide a general overview molecular diversity across the sites, rather than a precise estimate within each site.

## Results

We collected approximately 300 specimens across the four sites; however, our analysis focused on the 224 individuals that were successfully barcoded (Table 1; S1 Table). We were able to identify nearly 70% of individuals to genus level using the morphological approach, but species-level identification was only possible for 56 individuals (25%). Some of the most numerous taxa across sites included Leptophlebiidae (11.2 % of individuals), Libellulidae (11.2 %), Naucoridae (6.7 %), Notonectidae (6.3 %), Chironomidae (5.4%), Gerridae (4.9 %), Hydropsychidae (4.0 %), Perlidae (3.6 %), and Baetidae (3.6 %).

**Table 1.**
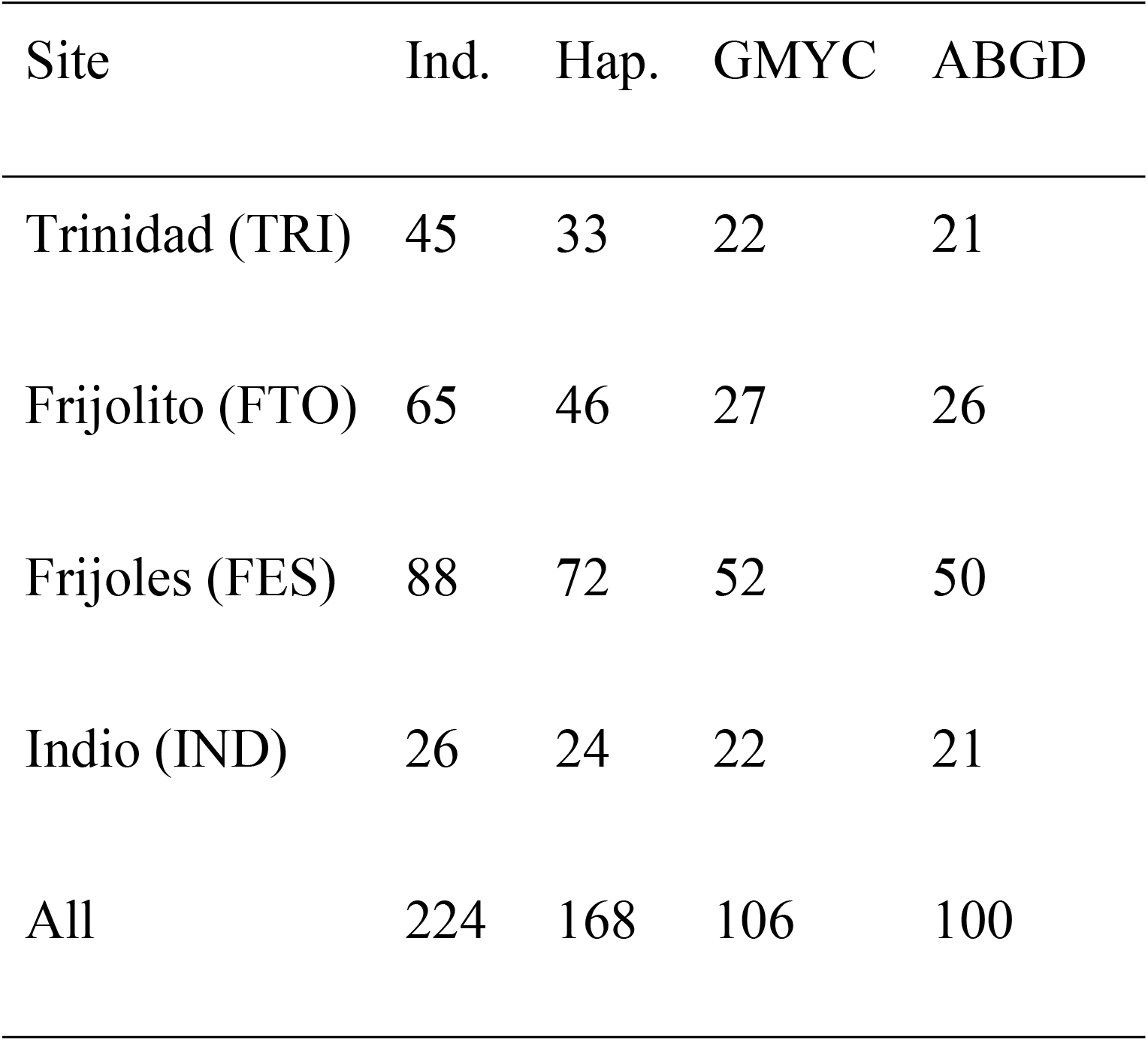
Macroinvertebrate lineage diversity in the Panama Canal Watershed.

The 224 COI sequences revealed a total of 168 haplotypes (S1 Table). After comparing these haplotypes with molecular data from Múrria et al [26], we found 153 (~91%) unique haplotypes from the Panama Canal Watershed. Of these haplotypes, only 8 had sequence lengths <620 bp (551-616 bp). The newly generated sequences were deposited in GenBank under accession numbers KX039451 - KX039650 and KU980966 - KU981004.

Our final COI dataset consisted of 171 terminals, including the 168 new barcode sequences, and 3 outgroup sequences retrieved from GenBank. The final aligned and pruned dataset contained 620 aligned positions, including gaps, with 371 variable sites, of which 359 were parsimony-informative (~96% of variable positions). As expected, we observed a hierarchical increase in the mean K2P genetic divergence with increasing taxonomic levels from within a species 0.38% (SE = 0.002), to within family 9.92% (SE = 0.01), to within order 19.75% (SE = 0.01). However, we were not able to identify our specimens to species level from our BLAST search, given that only around 50% of our sequences matched existing specimens in the public databases, and most of these matches conformed at the genus and family level only. Both ML and BI inference trees for all specimens showed well-defined clades at the level of order and family, with some differences in the topology, but overall support was higher for the BI tree, which we used to represent the number of molecular species (Fig 2; S1 Fig).

Our species delimitation analyses yielded variable, but relatively high numbers of species. Specifically, GMYC detected 106 MOTUs (95% confidence intervals: 104-109), whereas ABGD found a total of 100 MOTUs (Fig 2). These ABGD results were confirmed independently of the chosen model (Jukes-Cantor and Kimura), and were unaffected by changes of prior limits for intraspecific variation and threshold.

**Fig 2.**
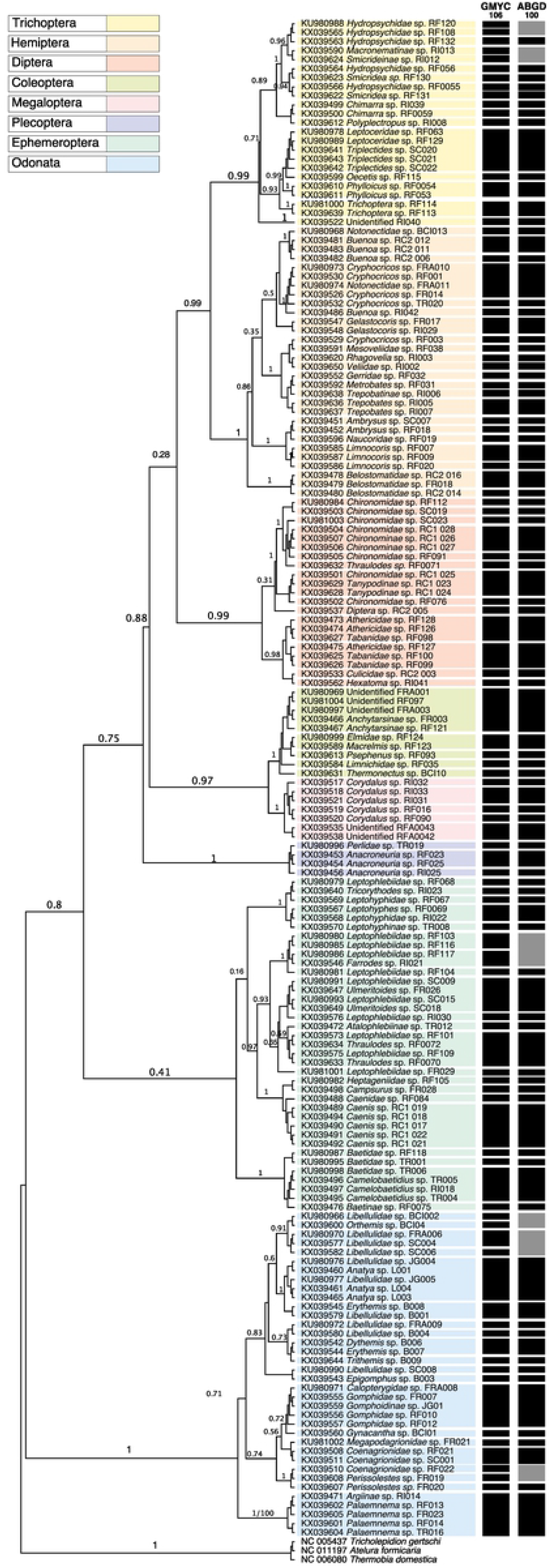
Molecular diversity in freshwater macroinvertebrates from Central Panama. The Bayesian inference tree shows species delimitation analyses based on generalized mixed Yule coalescent (GMYC) and the automatic barcode gap discovery (ABGD). Black and grey blocks represent putative molecular species, with taxa sharing the same block corresponding to similar species. Numbers next to the nodes represent Bayesian posterior probability values.

When looking at spatial patterns of diversity, we observed some overlap in the number of shared MOTUs as well as a considerable number of unique haplotypes in each river: Frijoles (65), Frijolito (43), Trinidad (31) and Indio (22) (Fig 3). This pattern was also supported by the Fisher’s alpha diversity index, which showed variation in molecular species among sites: Frijoles 53.40, Frijolito 22.80, Trinidad 17.01, and Indio 67.63. Whittaker’s index of β diversity also showed high species turn-over across sites (0.87). Similarly, our rarefaction/extrapolation analyses showed variation in species richness across sites: Frijoles (52), Frijolito (27), Trinidad (22) and Indio (22). However, the most striking pattern was a lack of saturation in the accumulation curves (Fig 4), and this pattern was consistent across the first three Hill numbers (Fig S2). Similar results were found when looking at diversity Hill across the Isthmus of Panama using the compiled barcoding data set. In particular, we observed substantial diversity of both MOTUs and unique haplotypes across sites (Fig 4), but the accumulation curves did not reach saturation (Fig 4). In addition, both MOTUs and haplotype diversity tended to increase at sites within the PCW, in contrast to sites located at the eastern and western portion of the country (Fig 4). Data on assignment and diversity of MOTUS across study sites are available in the supplementary material (S1 Table).

**Fig 3.**
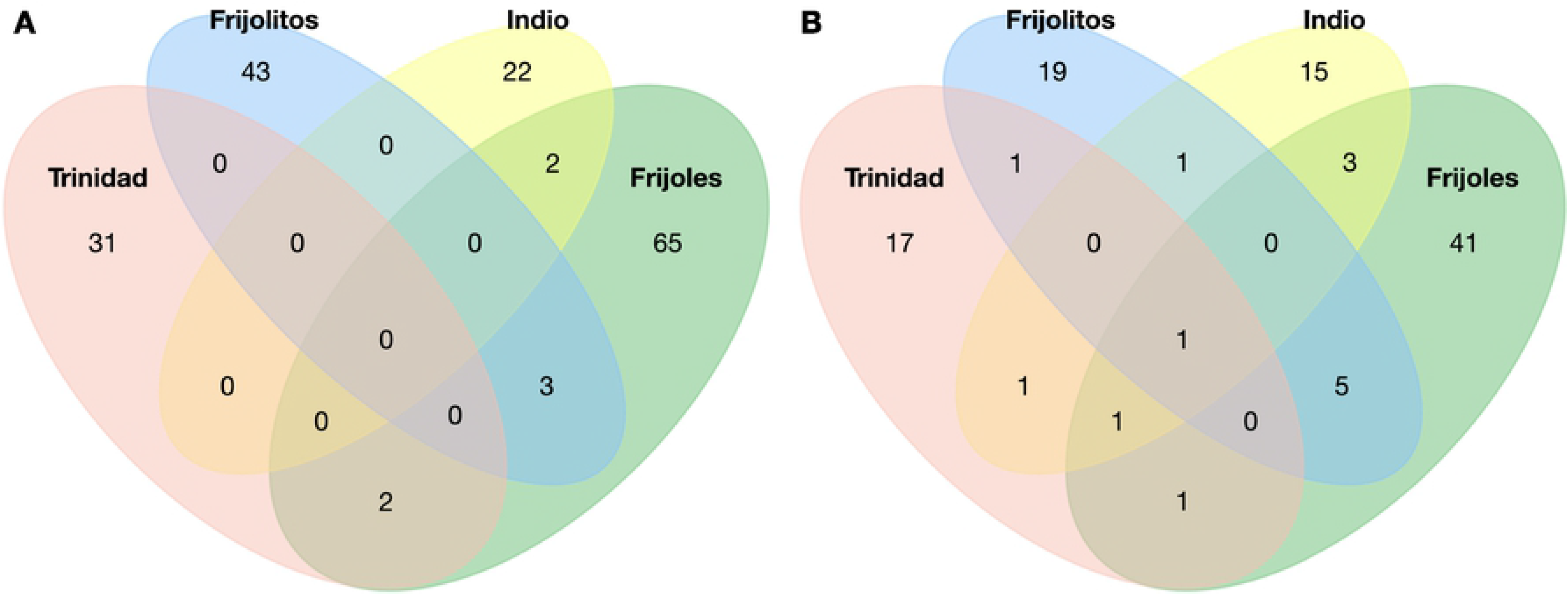
Distribution of molecular diversity in freshwater macroinvertebrates among streams within Panama Canal Watershed. Venn diagrams show the number of shared and unique haplotypes (A) and MOTUs (B) across four streams: Trinidad (pink), Frijolito (blue), Indio (yellow) and Frijoles (green).

**Fig 4.**
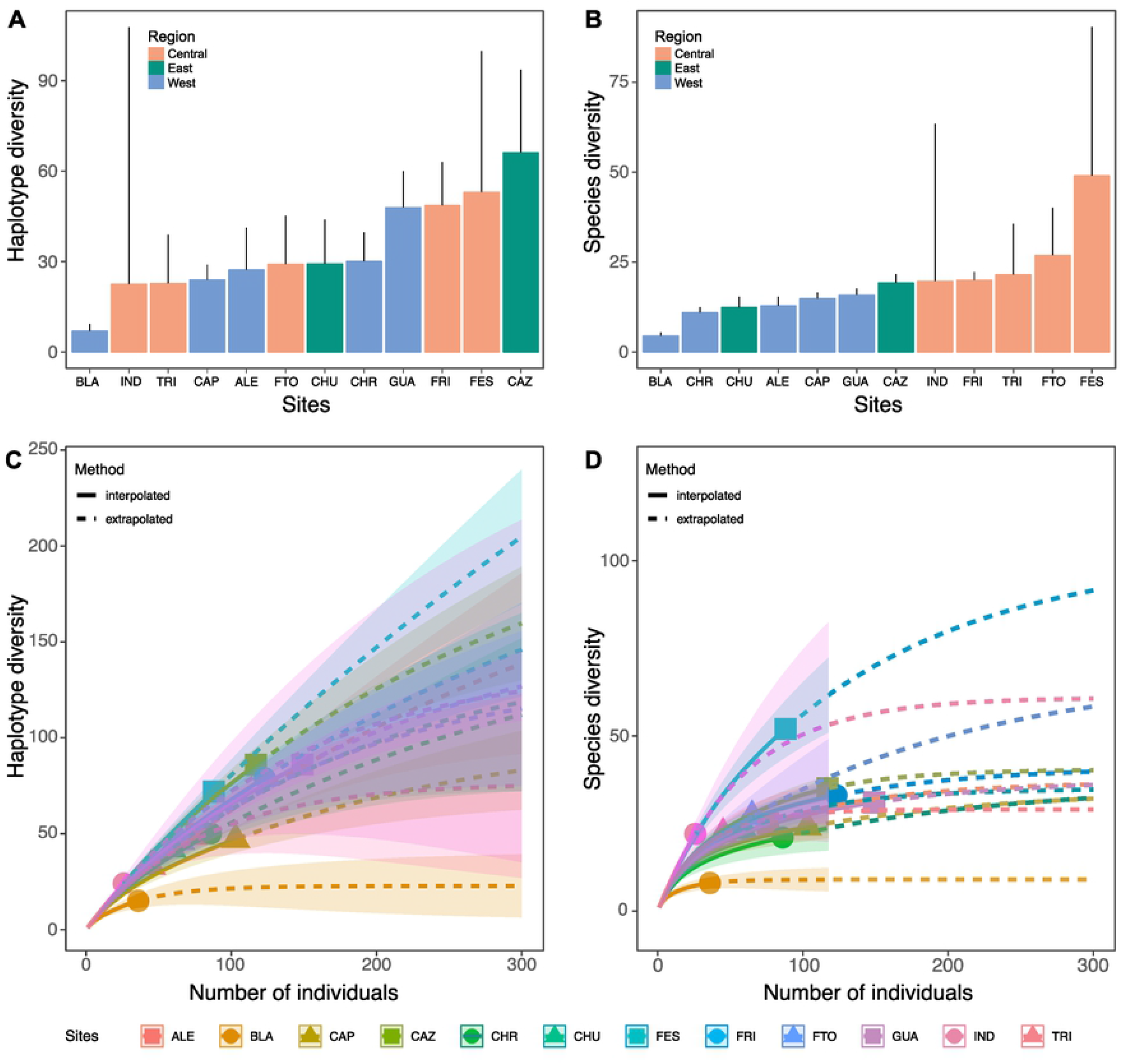
Molecular diversity in freshwater macroinvertebrates along the Isthmus of Panama. Panels show Simpson’s diversity index for both haplotype (A) and MOTUs (B) diversity across sites, and rarefaction and extrapolation curves for both haplotypes and MOTUs diversity each site. Sites are: Alemán (ALE), Chorro (CHO), Blanco (BLA), Guabal (GUA), Capira (CAP), Frijolitos (FRI), Cerro Azul (CAZ), and Chucanti (CHR) from Múrria et al. 2015; and Trinidad (TRI), Frijolitos (FTO), Frijoles (FES) and Indio (IND) (from present study). The Central region corresponds to the Panama Canal Watershed.

## Discussion

Using DNA-barcoding, we examined the diversity of freshwater macroinvertebrates at a small spatial scale, among streams within the Panama Canal watershed (PCW). We also compiled existing barcoding data [26] to contrast macroinvertebrate diversity at a broader spatial scale, across sites along the Isthmus of Panama. Overall, we found high lineage diversity across sites within the PCW (Table 1; Fig 2), and a large portion of these lineages appear to be unique to each site (Fig 3). In addition, our rarefaction/extrapolation approach showed that this diversity is still undersampled across sites both within the PCW and along the Isthmus of Panama (Fig 4).

These findings confirm that the diversity of freshwater macroinvertebrates in Neotropical environments is largely understudied [10,34,68], and could be much higher than previously thought. In addition, the fact that only a small number of specimens from these taxa matched available sequences in public databases further highlights the potential for biodiversity discovery in Neotropical freshwater environments. This seems particularly relevant for taxa such as Hydropsychidae, Gerridae, Chironomidae, Leptophlebiidae, Libellulidae and Notonectidae, which showed high lineage/haplotype diversity across sites (Table 1; Fig 2). Some of these taxa also showed high haplotype diversity in a previous molecular study across Panama [26], and are thought to hold an unknown number of undescribed species in the Central American Isthmus [6,10]. They also span several functional feeding groups such as filterers (Hydropsychidae), scrapers (Chironomidae) and predators (Notonectidae), which play a crucial role in freshwater ecosystems [1,2].

Contrary to our expectation, our results revealed high species turn-over at a small geographical scale. These findings complement recent work showing high haplotype endemicity among isolated watersheds across the Isthmus of Panama [26]. However, we expand on this work by highlighting the possibility that endemicity of Neotropical macroinvertebrates can be substantial even within a single watershed. In fact, only 15 of the 426 unique haplotypes reported by Múrria et al. [26] overlapped with haplotypes found here. Typically, diversification in freshwater organisms is marked by a strong geographic signature, where genetic divergence is facilitated by spatial isolation among populations [27,28]. However, the contribution of geographic isolation to diversification of Neotropical freshwater macroinvertebrates has received little attention to date [26]. In addition, the fact that most macroinvertebrates are semiaquatic, and are likely to disperse during the adult stages [16,17] may limit genetic isolation among nearby stream communities. However, our finding of high endemicity within the dispersal range of macroinvertebrates (e.g., Frijoles and Frijolitos are only separated by ~300m) suggests that Neotropical streams and rivers can be considered as continental islands of diversity.

Another interesting finding was that macroinvertebrate diversity appeared to increase at sites located in Central Panama, specifically within the PCW (e.g., Frijoles, Frijolito, Trinidad). Although it is unclear what may be driving this pattern, we suggest that increased dispersal and biodiversity exchange mediated by the construction of the Panama Canal and the rupture of the continental divide may be promoting the increase in local macroinvertebrate diversity. This process has been previously established for freshwater fishes [36,69], and it is very likely to be affecting other components of freshwater biodiversity, including macroinvertebrates. Another possibility is that conservation efforts around the PCW are facilitating persistence of freshwater biodiversity [70], in contrast with the eastern and western regions of the country, but additional research is needed to assess this possibility.

Our finding of high endemicity at small geographic scale is also relevant in the face of increasing anthropogenic disturbances [32,33,71–73]. Specifically, it suggests that small-scale local disturbances could have drastic consequences for the maintenance of a unique freshwater biodiversity - but this diversity is still unknown. We therefore predict that the current rate of species loss in freshwater ecosystems might be surpassing the rate of species discovery in Neotropical environments. Overall, however, further work is clearly needed to disentangle the contribution of other factors such as genetic drift, local adaptation, and environmental disturbance to persistence and diversification of Neotropical freshwater macroinvertebrates.

Taken together our results confirm the expectation that the diversity of Neotropical macroinvertebrates remains understudied. They also indicate that uncovering this hidden diversity is crucial to our understanding of the local and regional processes that shape biodiversity in Neotropical freshwater environments.

## Acknowledgments

We dedicate this study to our friend Ruth G. Reina. Her passion for tropical freshwater biology served as an inspiration to this work. Logistical support was provided by the Smithsonian Tropical Research Institute. Field assistance was provided by Celestino Martínez, Nohelys Alvarado, Carlos Nieto, and Débora Delgado. Diana Sharpe provided valuable comments on an earlier version of the manuscript. Celestino Aguilar is supported by the Sistema Nacional de Investigación (SNI). Luis F. De León is supported by the University of Massachusetts Boston.

## Supporting information

**S1 Fig. Rarefaction and extrapolation curves for molecular diversity (MOTUs) at each site**. Number at the top represent fit curves for the first three Hill numbers: species richness (q = 0), the exponential of Shannon entropy (“Shannon diversity”, q = 1), and the inverse Simpson concentration (“Simpson diversity”, q = 2), using individual-based abundance data. Sites are: Alemán (ALE), Chorro (CHO), Blanco (BLA), Guabal (GUA), Capira (CAP), Frijolitos (FRI), Cerro Azul (CAZ), and Chucanti (CHR) from Múrria et al. 2015; and Trinidad (TRI), Frijolitos (FTO), Frijoles (FES) and Indio (IND) (from present study).

**S2 Fig. Phylogenetic tree determined by the Maximum Likelihood (ML)**. Data represent *cox1* sequences obtained from 224 freshwater macroinvertebrates collected within the Panama Canal Watershed. Number on the tree branches show nodal support.

**S1 Table. Molecular and taxonomic diversity of freshwater macroinvertebrates within the Panama Canal Watershed**. For each specimen we show taxonomic group (i.e., order, family and genus), molecular species identity (MOTUs: based on ABGD and GMYC), Haplotype identity, Genbank accession number and sampling site. Sites are Trinidad (TRI), Frijolitos (FTO), Frijoles (FES) and Indio (IND).

